# Diversification of molecular pattern recognition in bacterial NLR-like proteins

**DOI:** 10.1101/2024.01.31.578182

**Authors:** Nathalie Béchon, Nitzan Tal, Avigail Stokar-Avihail, Sarah Melamed, Gil Amitai, Rotem Sorek

## Abstract

Antiviral STANDs (Avs) are bacterial anti-phage proteins that are evolutionarily related to immune pattern recognition receptors of the NLR family. Following recognition of a conserved phage protein, Avs proteins exhibit cellular toxicity and abort phage propagation by killing the infected cell. Type 2 Avs proteins (Avs2) were suggested to recognize the large terminase subunit of the phage as a signature of phage infection. Here, we show that while Avs2 from *Klebsiella pneumoniae* (KpAvs2) can be activated when heterologously co-expressed with the terminase of phage SECphi18, during infection *in vivo* KpAvs2 recognizes a different phage protein, named KpAvs2-stimulating protein 1 (Ksap1). We show that KpAvs2 directly binds Ksap1 to become activated, and that phages mutated in Ksap1 can escape KpAvs2 defense despite encoding an intact terminase. Our results exemplify the evolutionary diversification of molecular pattern recognition in bacterial Avs2 proteins, and highlight that pattern recognition during infection can differ from results obtained using heterologous co-expression assays.

## Introduction

The ability to detect pathogen invasion is a hallmark of immune systems across all domains of life^1–6^. Pattern recognition receptors, which detect molecular signatures indicative of pathogen infection, are at the front line of innate immune systems of many organisms^1,5–7^. These receptors can detect pathogen-associated molecular patterns (PAMPs) such as foreign nucleic acids^8^, conserved proteins^1,7^, and conserved non-proteinaceous molecules of the pathogen^1^.

The STAND NTPase superfamily of proteins is a large family of immune pattern recognition receptors present in animals, plants, fungi, archaea and bacteria^9–12^. These proteins typically have a tripartite architecture, consisting of a long C-terminal domain that senses the invader PAMP molecule, a central STAND NTPase domain of the NACHT or NB-ARC family, and an N-terminal domain that mediates cell death or an inflammatory response once an invader has been recognized^1,13,14^. The STAND NTPase superfamily includes animal inflammasomes and plant resistosomes, which are collectively named nucleotide-binding leucine-rich repeat containing receptors (NLRs) because their C-terminal sensor domains typically comprise leucine-rich repeats^9,15^.

In bacteria, STAND NTPase proteins were broadly found to protect against phage infection^7,10,12,16,17^. These proteins were dubbed NLR-like proteins^16,17^, bacterial NACHT (bNACHT)^12^, antiviral ATPase/NTPase of the STAND superfamily (AVAST), or antiviral STAND (Avs)^7,10^. Bacterial STAND NTPases preserve the tripartite structure of proteins in this family, and their N-terminal domains usually function as direct effectors of cell death^7,12^. Prokaryotic STAND NTPases were detected in 4%-10% of all published bacterial genomes^7,12^, and are thought to be the evolutionary ancestors of eukaryotic NLRs^9,12^.

A recent study has characterized a large family of Avs proteins in bacteria that were documented to recognize the large subunit of the phage terminase^7^, a highly conserved protein essential for phage DNA packaging^18^. Three terminase-recognizing Avs families were identified (Avs1, Avs2 and Avs3), and a cryo-EM structure of Avs3 from *Salmonella enterica* complexed with the phage terminase explained the structural basis for Avs3-terminase molecular recognition^7^. Co-expression of phage large terminase proteins with protein representatives from the Avs1, Avs2 and Avs3 families caused cellular toxicity, indicating that recognition of the terminase by these Avs proteins activates their toxic N-terminal effectors^7^.

In this study, we examined a defense system from *Klebsiella pneumoniae* that includes an Avs2-family protein. We show that, once it recognizes phage infection, the Avs2 protein indiscriminately degrades DNA to abort phage infection. Surprisingly, while Avs2 can be activated when heterologously co-expressed with the phage large terminase subunit as was previously reported, our data show that during infection, the *K. pneumoniae* Avs2 is activated by binding a phage protein of unknown function, here called Ksap1. We demonstrate direct interaction between the C-terminus of Avs2 and Ksap1, and show that phages in which Ksap1 is deleted or mutated become resistant to the Avs2 system of *K. pneumoniae*. Our findings expand the repertoire of known phage proteins recognized by bacterial NLR-like Avs proteins and underscore how evolutionary diversification allows these proteins to adapt to a range of phage targets.

## Results

### An Avs2-containing operon that protects *E. coli* against phage

A previous study identified over 2000 homologs of Avs2 in various bacterial genomes^7^. We became interested in a set of homologs typified by Avs2 from *Klebsiella pneumoniae* S_15PV (KpAvs2) that seems to be embedded in a three-gene operon, as opposed to typical Avs2 proteins that usually function as a single protein (Figure 1A and Supplementary Table S1). This operon was frequently encoded in proximity to other known defense systems in microbial genomes, supporting the notion that its role is to defend against phages (Figure 1B). As expected, HHpred^19,20^ analysis showed that KpAvs2 is divided into three distinct domains: an N-terminal endonuclease domain of the Mrr restriction endonuclease family, a central STAND NTPase domain, and a long C-terminal domain which displayed sequence homology to the C-terminal domains of previously characterized Avs2 proteins^7^. The two associated genes encoded a protein with a radical S-adenosylmethionine (SAM) domain, and a protein of unknown function with an HHpred^19,20^ hit to the pfam protein family PF19902 (domain of unknown function DUF6375) (Figure 1A). We denote these proteins Avs-associated protein 1 (Avap1) and Avs-associated protein 2 (Avap2).

**Figure 1.**
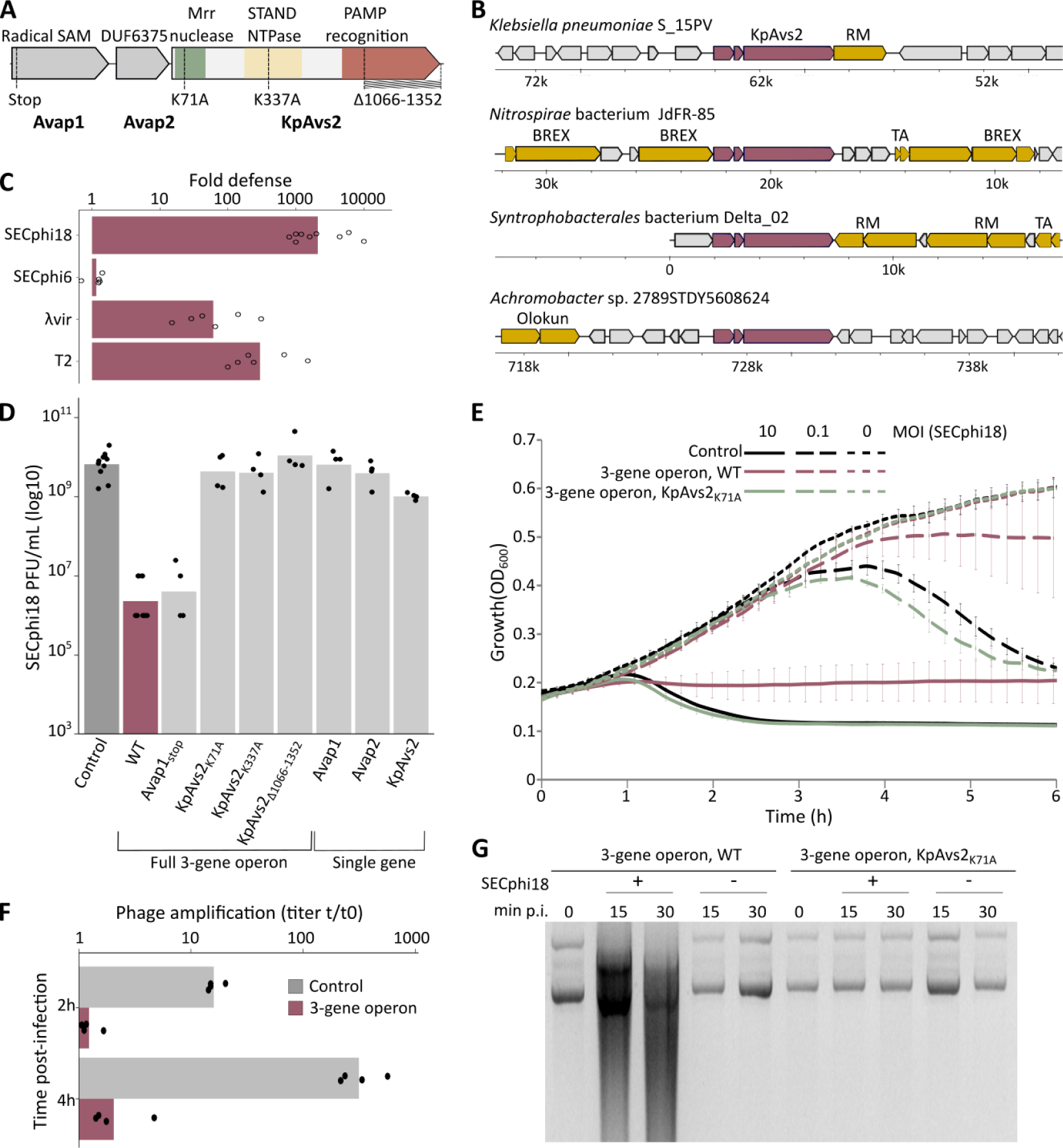
The KpAvs2 operon protects against phage by abortive infection. **A.** Domain organization of the KpAvs2 operon from *Klebsiella pneumoniae* S_15PV. Mutations used in this study are indicated below the operon. **B.** Gene neighborhoods of selected KpAvs2 homologs. Purple, homologs of the KpAvs2 operon; orange, known defense genes. Presented genomes are *Klebsiella pneumoniae* S_15PV (IMG^22^ scaffold identifier ID: 2701097382), *Nitrospirae* bacterium JdFR-85 (scaffold ID: 2728441048), *Syntrophobacterales* bacterium Delta_02 (scaffold ID: 2751221707), *Achromobacter* sp. 2789STDY5608624 (scaffold ID: 2660299158). RM: Restriction-modification, BREX: Bacteriophage Exclusion, TA: toxin-antitoxin. **C.** Fold defense, calculated as the ratio between the efficiency of plating of phages infecting control *E. coli* cells and cells expressing the KpAvs2 operon. Experiments were performed at 25°C for phages SECphi18 and SECphi6, or 37°C for λvir and T2. Bar graph represents average of 5-9 independent replicates, with individual data points overlaid. **D.** Efficiency of plating of SECphi18 phages infecting *E. coli* cells expressing the KpAvs2 three-gene operon with the indicated mutations, infected at 25°C. Data represent plaque-forming units (PFU) per mL. Bar graph represents average of 4 independent replicates for each mutant, and 11 replicates for both negative and positive control strains, with individual data points overlaid. Negative control is a strain in which GFP is expressed instead of the defense system. **E.** Liquid culture growth of *E. coli* cells expressing the WT KpAvs2 operon, an operon with a KpAvs2 mutated in the nuclease effector domain, or control cells expressing GFP. Cells were infected by phage SECphi18 at 25°C. Each curve represents the average of three replicates, error bars represent standard deviation. MOI: multiplicity of infection. **F.** Titer of SECphi18 phage propagated on *E. coli* cells expressing either the KpAvs2 operon or GFP as a control. Data represent the titer of SECphi18 measured in PFU/mL after two or four hours from initial infection, corresponding roughly to one and two cycles of infection, divided by the original phage titer prior to infection. Infection was performed at MOI=0.1. Bar graph represents average of 4 replicates, with individual data points overlaid. **G.** Ethidium bromide-stained agarose gel of plasmids extracted from infected cells expressing either the WT KpAvs2 operon or an operon in which KpAvs2 was mutated in the nuclease domain. DNA was extracted at 0, 15 and 30 min post infection (min p.i.) by phage SECphi18 at MOI=10 and 25°C. As a control, plasmids were extracted from *E. coli* cultures not challenged by phages.

To test whether the KpAvs2-containing operon is able to protect against phage infection, we heterologously expressed it in *Escherichia coli* using an arabinose-inducible promoter. Plaque assay experiments showed that cells expressing this operon became resistant to multiple phages, with the strongest protection observed against SECphi18, a phage from the *Dhillonvirus* genus (Figure 1C and Supplementary Figure S1). A point mutation predicted to disrupt the active site of the N-terminal endonuclease domain of KpAvs2, K71A, abolished defense, confirming that the nuclease activity is essential for defense as expected from previous studies on Avs proteins^7,12^ (Figure 1D and Supplementary Figure S2AB). Similarly, a point mutation disrupting the STAND NTPase active site, K337A, and a deletion of the C-terminal PAMP recognition domain, also abolished defense (Figure 1D and Supplementary Figure S2AB).

Expression of KpAvs2 alone conferred only weak defense against phages, suggesting that one or both of the associated genes are necessary for the full defense capacity; and neither Avap1 nor Avap2 conferred defense when expressed alone (Figure 1D and Supplementary Figure S2C). Expressing the three-gene operon in which *avap1* was inactivated by a premature stop codon protected the culture from SECphi18 infection as efficiently as the WT three-gene operon, suggesting that Avap1 is dispensable for defense against SECphi18 (Figure 1D and Supplementary Figure S2AB). In contrast, inactivation of Avap1 did reduce defense against phage T2, suggesting that Avap1 is required for defense against some, but not all, phages (Supplementary Figure S3). Attempts to delete Avap2 from the operon were unsuccessful, implying that perturbing this gene in the presence of Avap1 and KpAvs2 might lead to cellular toxicity.

Given that KpAvs2 protected against SECphi18 only in the presence of Avap2 (Figure 1D), we investigated the possibility of an interaction between these two proteins. AlphaFold-Multimer^21^ analysis predicted high confidence interactions between Avap2 and an alpha-helical bundle extending between the nuclease domain and the STAND NTPase domain of KpAvs2 (Supplementary Figure S4A, S5A). Co-immunoprecipitation experiments confirmed physical binding between Avap2 and KpAvs2 (Supplementary Figure S4BC). Notably, Avap2 is not predicted to interact with the distal part of the C-terminal domain within KpAvs2 that likely comprises the phage recognition pocket (see below) (Supplementary Figure S4A). In agreement with the AlphaFold-Multimer prediction, Avap2 retained its ability to bind KpAvs2 even when this C-terminal domain was deleted from KpAvs2 (Supplementary Figure S4BC). These results show that KpAvs2 binds the accessory protein Avap2, suggesting that this binding may contribute to the defensive function.

Infection experiments in liquid culture showed that the KpAvs2 operon protected the culture when cells were infected by phage SECphi18 at a low multiplicity of infection (MOI = 0.1). Infection at a high MOI resulted in growth arrest of cells expressing the KpAvs2 operon, occurring earlier than the time in which control cells lysed following infection by SECphi18 (Figure 1E). Consistently, plaque-forming units (PFU) analysis showed that phages were unable to replicate on KpAvs2-expressing cells (Figure 1F), confirming that KpAvs2 protects via abortive infection.

Previous studies with Avs3 and Avs4 have shown that these proteins are phage-activated DNA endonucleases that non-specifically degrade DNA upon phage recognition^7^. As KpAvs2 encodes an N-terminal endonuclease domain, we hypothesized that this protein, too, would degrade DNA in response to phage infection. In agreement with this hypothesis, analysis of DNA extracted from cells infected by phage SECphi18 showed a distinct DNA smear pattern indicative of non-specific DNA degradation in cells expressing KpAvs2 (Figure 1G). Collectively, our findings identify an Avs2 variant that necessitates an accessory protein and non-specifically degrades DNA as a defense mechanism to prevent phage replication.

### KpAvs2 is activated *in vivo* by a small phage protein of unknown function

Avs2 proteins were previously shown to become toxic when co-expressed with phage large terminase subunit proteins, and it was therefore hypothesized that the large terminase is the PAMP sensed by Avs2^7^. To test if the toxic effects of KpAvs2 can be activated by the phage terminase, we attempted to transform plasmids encoding the large terminase subunit of phage SECphi18 into cells that encode KpAvs2. We observed a substantial reduction in transformation efficiency of a plasmid encoding the SECphi18 terminase in the presence of WT KpAvs2 compared to a KpAvs2 mutant lacking the C-terminal domain (Figure 2A). These findings suggest that the large terminase subunit of SECphi18 is toxic in the presence of KpAvs2, which led us to initially think that this phage protein may serve as the activator of KpAvs2.

**Figure 2:**
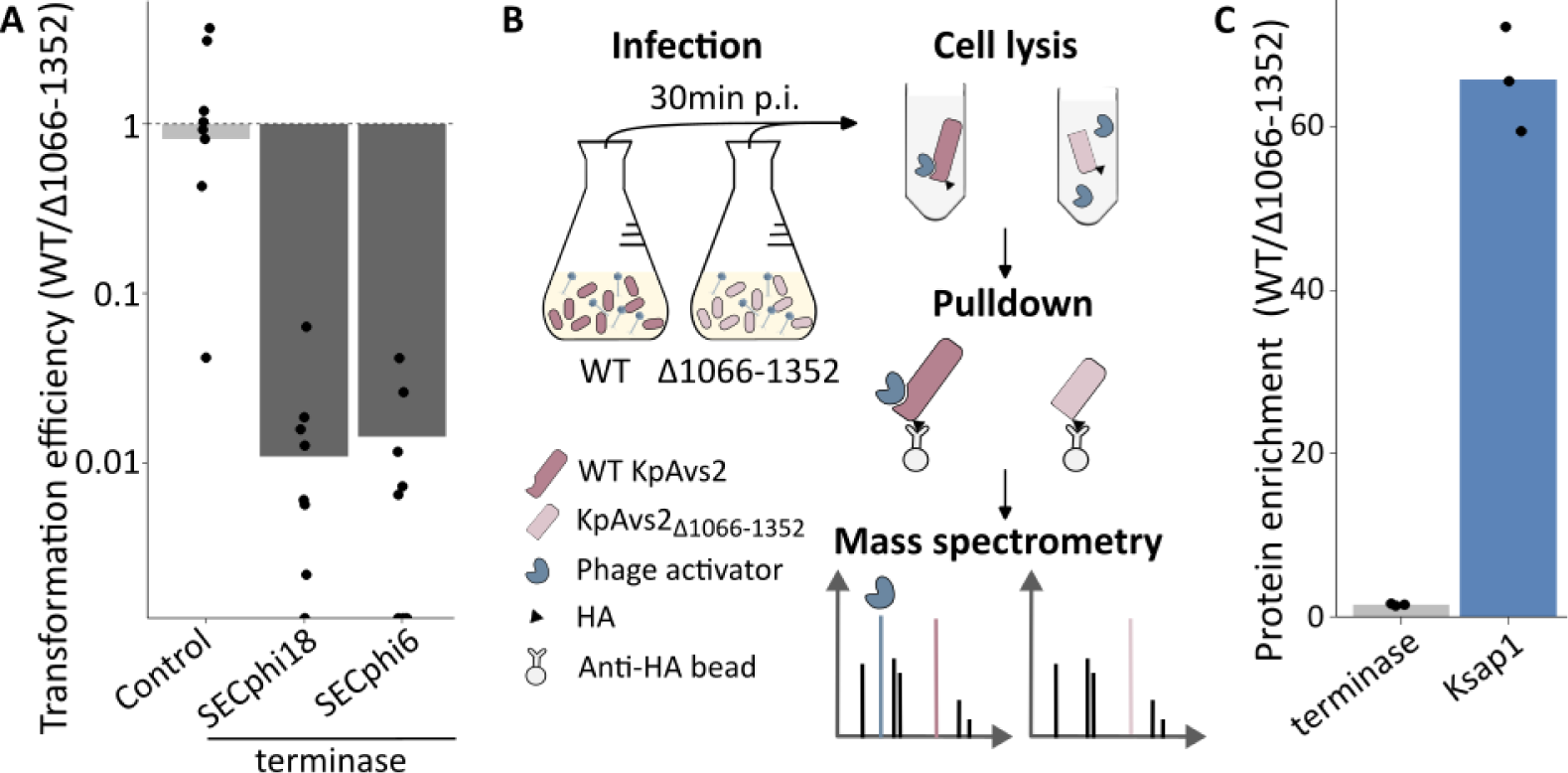
KpAvs2 recognizes SECphi18 large terminase subunit in toxicity assays but not during infection. **A.** Transformation efficiency of the gene encoding the large terminase subunit of SECphi18 or SECphi6. Data represent the ratio of transformants obtained using bacteria encoding the WT KpAvs2 operon divided by transformants obtained with the KpAvs2_Δ1066-1352_ deletion. Bar graph represents average of 8 replicates, with individual data points overlaid. Control indicates transformation with a plasmid encoding RFP instead of the terminase. **B.** Schematic of the protein pulldown experiment. Cultures expressing HA-tagged KpAvs2 or KpAvs2_Δ1066-1352_ were infected by phage SECphi18 at MOI of 5. At 30 min post infection, co-immunoprecipitation using anti-HA antibodies was performed on cell lysates, and pulled-down proteins were analyzed by mass spectrometry to identify proteins that were enriched when WT KpAvs2 was used as bait as compared to the mutated KpAvs2. **C.** Mass spectrometry analysis of SECphi18 proteins pulled down with KpAvs2 during infection. Presented are data for the SECphi18 large terminase subunit and Ksap1, a list of all identified proteins is in Supplementary Table S2. Data presented as the ratio between protein abundance in the WT KpAvs2 sample and the KpAvs2_Δ1066-1352_ sample. Protein abundance was normalized based on bait abundance for each sample. Average of 3 replicates, individual data points overlaid.

If KpAvs2 indeed recognizes the phage large terminase subunit as a trigger for infection, one would expect KpAvs2 to bind this phage protein during infection. To test whether this is the case, we infected cells expressing an HA-tagged KpAvs2 with phage SECphi18, and immunoprecipitated the tagged KpAvs2 together with proteins bound to it. Surprisingly, mass spectrometry analysis of proteins that were pulled down together with KpAvs2 did not show enrichment for the phage large terminase subunit (Figure 2BC, Supplementary Table S2). Rather, another phage protein, which we denote here KpAvs2-stimulating protein 1 (Ksap1), was 60-fold enriched in the KpAvs2 pulled-down sample as compared to similar samples where the KpAvs2 C-terminal recognition domain was deleted (Figure 2C). These results implied that, counter to our original hypothesis, a phage protein other than the large terminase subunit binds KpAvs2 during infection.

Ksap1 is a 74 amino-acid long protein of unknown function that resides between a methyltransferase- and a phosphatase-encoding genes in the SECphi18 genome (Figure 3A). To test if the interactions observed during infection between KpAvs2 and Ksap1 could be reconstituted *in vitro*, we performed co-immunoprecipitation assays with a FLAG-tagged Ksap1 and an HA-tagged KpAvs2. To avoid possible toxicity, each of the proteins was expressed in a separate culture, and the proteins were mixed *in vitro* on beads. These co-immunoprecipitation assays demonstrated physical interactions between the two proteins, which were abolished when the C-terminus of KpAvs2 was deleted, confirming our observations from the pulldown assay during infection (Figure 3B). Structural modeling with AlphaFold-Multimer^21^ showed high-scoring predicted interactions between Ksap1 and the C-terminal domain of KpAvs2, offering a structural explanation for the binding between the two proteins (Figure 3C and Supplementary Figure S5B).

**Figure 3.**
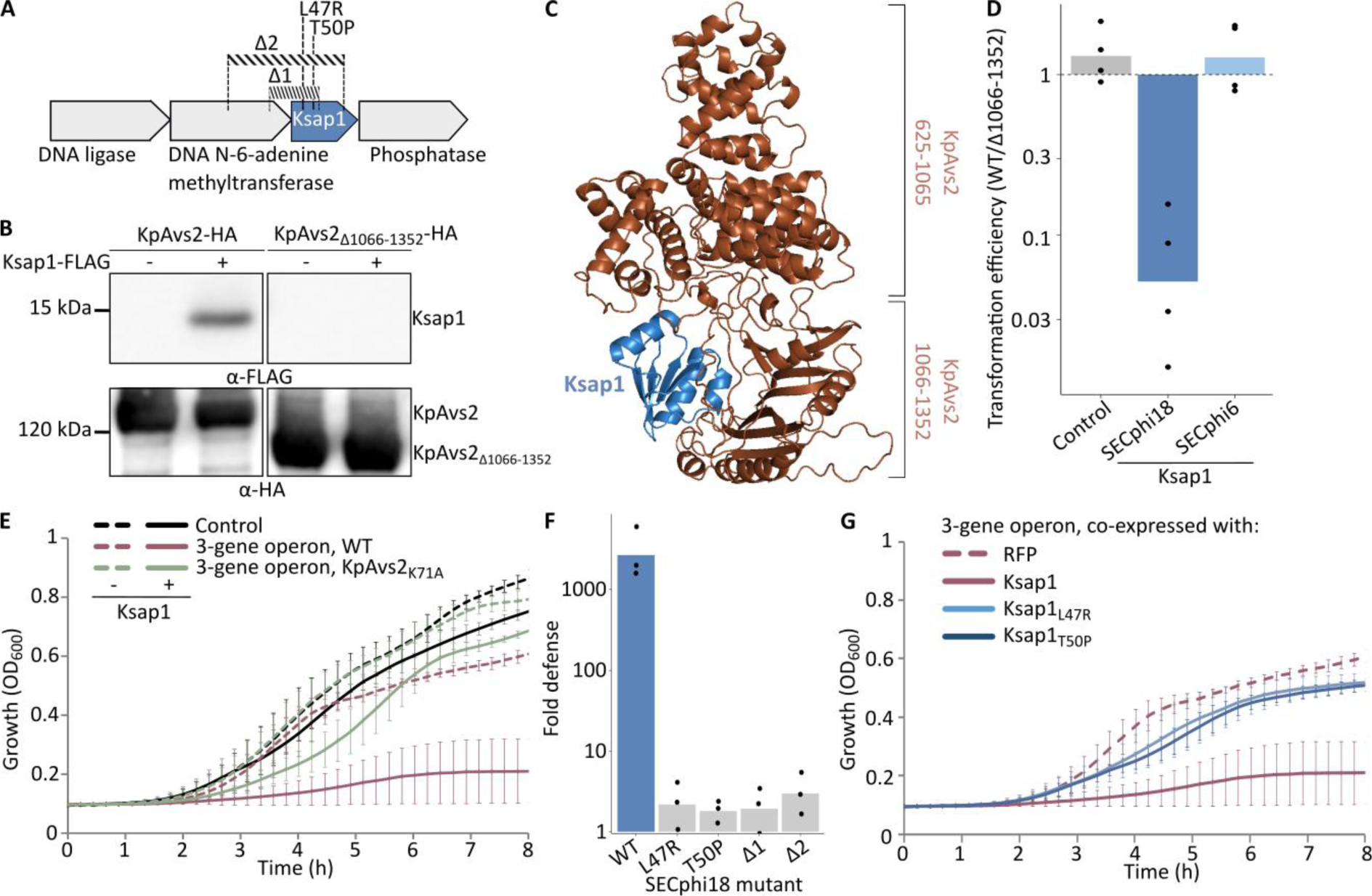
KpAvs2 defense is activated by direct binding to a small phage protein of unknown function. **A.** Genetic organization of SECphi18 genes surrounding *ksap1*. Mutations identified in escaper phages are represented on top. Δ1 and Δ2 represent deletions of part of the locus, L47R and T50P represent missense mutations at the indicated amino acids. **B.** Co-immunoprecipitation of HA-tagged KpAvs2 and 3xFLAG-tagged Ksap1. α-HA beads were used to immunoprecipitate HA-tagged KpAvs2 or KpAvs2_Δ1066-1352_, and the interacting Ksap1 was detected by western blot against FLAG tag. A western blot against HA tag was performed on the same blot to control for efficiency of pulldown (lower panel). **C.** AlphaFold-Multimer^21^ predicted interactions between Ksap1 (in blue) and the C-terminal domain of KpAvs2 (brown). Model confidence score: 0.84. **D.** Transformation efficiency of the gene encoding Ksap1 of SECphi18 or SECphi6. Data represent the ratio of transformants obtained using bacteria expressing the WT KpAvs2 operon divided by transformants obtained with the KpAvs2_Δ1066-1352_ deletion. Bar graph represents average of 4 replicates, with individual data points overlaid. **E.** Growth curves of *E. coli* cells co-expressing both the KpAvs2 operon and Ksap1. Cells expressed either the KpAvs2 operon or GFP (Control). Additionally, cells expressed either Ksap1 (+) or RFP as a control (-). Data represent the average of 6 replicates, error bars represent standard deviation. **F.** Fold defense, calculated as the ratio of the efficiency of plating of SECphi18 phages on *E. coli* control cells that express GFP and cells expressing the KpAvs2 system. Infection was performed at 25°C. Data presented for WT or mutated SECphi18 phages. Bar graph represents the average of 3 independent replicates, with individual data points overlaid. **G.** Growth curves of *E. coli* cells co-expressing the KpAvs2 operon and Ksap1 variants. Presented data are as in panel E.

To test whether KpAvs2 becomes toxic in the presence of Ksap1, we attempted to transform plasmids carrying *ksap1* into cells expressing the KpAvs2 operon. This transformation was substantially less efficient than transformation of a control plasmid (Figure 3D), and transformants frequently carried suppressor mutations, suggesting toxicity. Nevertheless, we managed to obtain a non-mutated transformant for further analysis. Expression of the KpAvs2 operon in the presence of Ksap1 resulted in cellular toxicity, which was abolished when KpAvs2 was mutated in its nuclease domain, confirming that Ksap1 activates KpAvs2 (Figure 3E).

Multiple recent studies have shown that examining phage mutants that escape specific defense systems can generate valuable insights into the mechanism of defense activation, because escaper phages can evade bacterial immunity by mutations in the genes that activate the bacterial defense system^23–27^. We propagated SECphi18 phages on *E. coli* expressing the KpAvs2 operon, and isolated six phage mutants that were able to escape defense. Notably, all six escaper phages harbored mutations in Ksap1 (Supplementary Table S3). The mutations included partial deletion of *ksap1* together with part of the upstream gene, or single point mutations altering leucine 47 or threonine 50 of Ksap1 to arginine and proline, respectively (Figure 3A and Supplementary Table S3).

We confirmed that bacteria expressing the KpAvs2 operon failed to defend against escaper phages carrying mutations in *ksap1* (Figure 3F). Co-expression of KpAvs2 with Ksap1_L47R_ and Ksap1_T50P_ did not cause growth arrest, suggesting that the Ksap1 variants in the mutated phages do not activate KpAvs2 and explaining why these phages escaped defense (Figure 3G and Supplementary Figure S6A). Notably, while one of the six escaper mutants carried a missense mutation in the gene encoding the large terminase in addition to a frameshift mutation in *ksap1*, the other five escaper phages were not mutated in the terminase, showing that phages could escape the KpAvs2 system defense even when expressing a wild-type terminase. These results further support the hypothesis that Ksap1, rather than the terminase, is the component sensed by KpAvs2 as a signature of SECphi18 infection.

### Natural variants of Ksap1 escape KpAvs2 defense

Phage SECphi18 is taxonomically closely related to phage SECphi6^28^. Both phages belong to the *Dhillonvirus* genus, and their ∼45 kb genomes show 92% sequence similarity over 96% of the sequence. Despite the similarity between the two phages, the KpAvs2 operon shows no defense against phage SECphi6 while strongly defending against SECphi18 (Figure 1C). SECphi6 harbors a homolog of Ksap1 in a locus syntenic to the Ksap1 locus in phage SECphi18, and the SECphi6 Ksap1 homolog differs from the SECphi18 Ksap1 homolog only by 14 residues (Figure 4A). We hypothesized that the altered sequence of the SECphi6 Ksap1 protein allows the phage to escape detection by KpAvs2. In support of this hypothesis, co-expression of KpAvs2 with Ksap1 from SECphi6 did not result in cellular toxicity (Figure 3D and 4B), suggesting that Ksap1 from this phage does not activate the KpAvs2 system.

**Figure 4:**
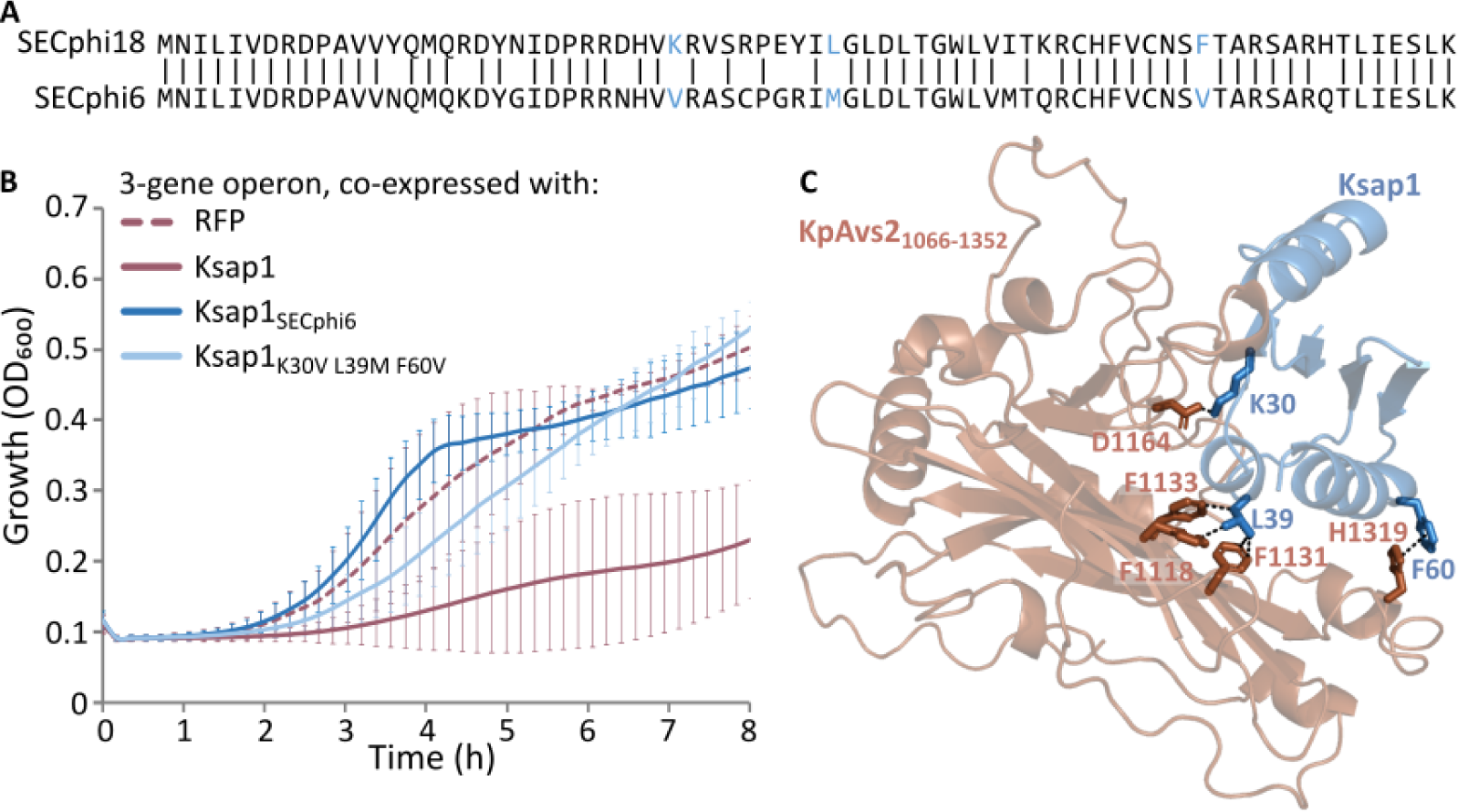
Natural variants of Ksap1 evade recognition by KpAvs2. **A.** Amino acid sequence alignment of Ksap1 proteins from SECphi18 and SECphi6 phages. Blue color indicates the amino acids that were mutated in panel B. **B.** Growth curves of *E. coli* cells co-expressing both the KpAvs2 operon and different variants of Ksap1. As controls, a plasmid expressing RFP instead of Ksap1 was used. Data represent the average of 6 replicates, error bars represent standard deviation. **C.** Interaction predicted between indicated residues of Ksap1 from SECphi18 (in blue) and KpAvs2 (brown). Interactions were predicted by AlphaFold-Multimer^21^ and RING^29^. Shown is a close-up of the structure presented in Figure 3C.

We used AlphaFold-Multimer^21^ to predict which amino acids are in direct interaction between Ksap1 and the C-terminal domain of KpAvs2. Among the 14 residues that differed between the SECphi18 Ksap1 and its homolog in SECphi6, three were predicted to make a strong interaction with KpAvs2. These were K30, generating an ionic bond; L39, generating van der Waals interactions with three separate residues in KpAvs2; and F60, generating a π-π stacking interaction (Figure 4C and Supplementary Table S4). These interactions were not present when we modeled the interactions of KpAvs2 with the SECphi6 version of Ksap1 (Supplementary Table S5), in which these residues are altered to valine, methionine and valine, respectively. We therefore hypothesized that the alteration of these three residues may explain why KpAvs2 does not protect from SECphi6. To examine this hypothesis, we replaced the three amino acids in the SECphi18 protein by their respective SECphi6 variants, and tested whether Ksap1_K30V L39M F60V_ could activate the toxicity of KpAvs2. Co-expression of Ksap1_K30V L39M F60V_ with the KpAvs2 operon was not toxic (Figure 4B and Supplementary Figure S6B), showing that alteration of these residues is sufficient for escape from KpAvs2 activation. Collectively, our results provide compelling evidence that Ksap1 functions as the PAMP that is directly recognized by KpAvs2 as a signature for SECphi18 infection, and that phages can escape KpAvs2 defense by altering Ksap1.

## Discussion

In this study, we characterized KpAvs2, a type 2 Avs system, which, akin to previously described type 2 Avs proteins^7^, operates through the recognition of a phage protein via physical binding. Recognition of the phage protein activates an N-terminal effector in KpAvs2, which non-specifically degrades cellular DNA. Our data confirm that the KpAvs2 operon becomes toxic when co-expressed with the large terminase subunit of the phage, as reported^7^. Such results were previously interpreted as if the PAMP naturally recognized by Avs2 is the large terminase protein of the phage. However, our data surprisingly show that the KpAvs2 operon recognizes a different protein when protecting against phage SECphi18 infection. We found that KpAvs2 binds Ksap1 during infection, and that this protein activates KpAvs2 defense. In support of these observations, the KpAvs2 system did not protect against phages mutated in Ksap1, although the large terminase subunit in these phages was intact. Moreover, KpAvs2 did not protect against SECphi6 at all, although the SECphi6 terminase seemed to be toxic in the presence of KpAvs2 based on our transformation efficiency assays (Figure 1C and 2A). These data collectively suggest that KpAvs2 recognizes Ksap1 as its PAMP, and not the phage terminase, when protecting against SECphi18. Future studies will be necessary to reveal why KpAvs2 is not activated by the phage terminase during infection.

Ksap1, the SECphi18-encoded activator of KpAvs2, is a small protein of unknown function. We were able to find homologs of this protein only in phages from the genus *Dhillonvirus,* where this protein is conserved in almost all sequenced phages of this genus (Supplementary Figure S7A and Supplementary Table S6). Sequence-based homology searches did not find significant homology to any protein of known function, and structure-based searches using Foldseek^30^ did not retrieve hits to any protein with an experimentally-determined structure. Foldseek search against a database of AlphaFold2^31^-generated protein structures (AFDB50^32^) did retrieve several marginal hits to proteins with a predicted response regulator domain, a protein domain that can be activated by phosphorylation and is typical to two-component signaling systems in bacteria^33^ (Supplementary Figure S7BC). However, a conserved aspartate residue that is essential for phosphorylation of response regulator domains is missing from Ksap1, suggesting that Ksap1 may not function as a response regulator (Supplementary Figure S7BC)^33^. The presence of Ksap1 in multiple *Dhillonvirus* phages suggests that it has a role in the biology of these phages, but since the phages could tolerate large deletions in the *ksap1* gene, we were not able to discern its functional role.

It is unclear how KpAvs2 can be activated, at least in co-expression assays, by both the large terminase and Ksap1. These two proteins show no sequence homology, and attempts to detect structural homology between these two proteins did not retrieve high-scoring alignments. These data suggest that Ksap1 and the large terminase are not evolutionarily related.

Contrasting with previously characterized type 2 Avs, for which a single protein is sufficient to provide defense, KpAvs2 necessitates at least one additional accessory protein. We found that the accessory protein Avap2 is essential for defense against SECphi18, and that this protein physically interacts with KpAvs2; however, its functional role is unclear. We also found that the other accessory protein, Avap1, is necessary for defense at least against some phages. Avap1 has a predicted radical SAM domain, similar to prokaryotic viperins (pVips) that generate antiviral compounds as a mode of defense^28^. We were unable to detect the modified nucleotides typical of pVip activity in cells expressing the full KpAvs2 operon during infection, and AlphaFold2^31^ prediction of Avap1 structure does not support the presence of a catalytic site that could accommodate an NTP, the substrate of pVips. Further studies will be necessary to decipher the roles of Avap1 and Avap2 in KpAvs2 defense.

Collectively, our findings underscore the evolutionary plasticity of PAMP recognition in defense systems. We extend the range of known phage proteins recognized by Avs systems by describing Ksap1 as the PAMP recognized by KpAvs2. Our results show that recognition data obtained based on co-expression experiments alone can, sometimes, be insufficient to discern PAMP identity during infection. This work extends our understanding of the diversification of molecular pattern recognition in a large family of immune receptors conserved across the tree of life.

## Materials and method

### Strains and growth conditions

*E. coli* K-12 MG1655 was grown in MMB media (lysogeny broth (LB) supplemented with 0.1 mM MnCl_2_ and 5 mM MgCl_2_) at 37°C or 25°C with 200 rpm shaking or on solid LB 1.5% agar plates. Ampicillin 100 μg/mL or kanamycin 50 μg/mL were added when necessary for plasmid maintenance. Induction was performed with 0.2% arabinose or 25 ng/mL anhydrotetracycline (aTc). Glucose 1% was used to repress the arabinose-inducible promoter when needed. All chemicals were obtained from Sigma Aldrich unless stated otherwise.

To propagate phages, wild type *E. coli* K-12 MG1655 was grown to OD_600_ ∼0.4-0.6 and infected with a sample from a single plaque of the phage then incubated until total culture lysis at 37°C, 200 rpm shaking. The lysate was then propagated again on a bacterial culture in the same way to increase phage titer. The lysate was then centrifuged 15 min at 3900 *g* and filtered through 0.2 μm filter and kept at 4°C until use. When necessary, phages were diluted in either MMB or phage buffer (50 mM Tris pH 7.4, 100 mM MgCl2, 10 mM NaCl).

A table of all plasmids, strains and phages used in this study can be found in Supplementary Table S7.

### Plasmid and strain construction

Target DNA was amplified using KAPA HiFi HotStart ReadyMix (Roche) according to manufacturer instruction or synthesized by Twist Bioscience (Supplementary Table S7). Supplementary Table S8 lists all primers used in this study. All primers were obtained from Sigma Aldrich.

Some plasmids were synthesized and cloned by GenScript Corporation as indicated in Supplementary Table S7. Others were cloned as described below.

For one-fragment DNA cloning, a linear plasmid obtained by PCR was ligated using KLD enzyme mix (NEB) for 5 min at room temperature before transformation into 5-alpha Competent *E. coli* High Efficiency (NEB) through heat shock. Briefly, competent cells were incubated with DNA for 30 min on ice then 30 s at 42°C before resuspension in MMB media. Cells were left to recover for 1h at 37°C, and were then plated on selective media.

For assembly of more than one fragment, PCR products were treated with FastDigest DpnI (Thermo Fisher Scientific) restriction enzyme for 30 min at 37°C to remove all remaining circular template plasmid and the enzyme was inactivated for 20 min at 80°C. The fragments were then assembled using NEBuilder HiFi DNA Assembly Master Mix (NEB) at 50°C for 30 min then transformed in 5-alpha Competent *E. coli* High Efficiency (NEB), as described above.

Colonies were checked by PCR using DreamTaq Green PCR Master Mix (Thermo Fisher Scientific) then Sanger sequenced through the DNA Sequencing unit of the Life Sciences core facilities of Weizmann Institute of Science; alternatively, plasmids were prepared using QIAprep Spin Miniprep Kit (Qiagen) according to manufacturer instructions and sent for whole plasmid sequencing at Plasmidsaurus (www.plasmidsaurus.com). Sequence-verified colonies were then transformed into electrocompetent *E. coli* MG1655 cells or into the relevant background and kept in 20% glycerol at -80°C.

### Plaque assays

Phage infectivity was assessed using the small drops plaque assay as described previously^34^. Briefly, 300 μL of overnight culture was mixed with 30 mL melted 0.5% agar MMB media containing the relevant inducers and poured into a 10 cm square petri dish, then left to recover for ∼1h at room temperature to allow expression of the induced proteins. 10 μL drops of 10-fold serial dilutions of phages were then dropped over the lawn of bacteria, allowed to dry, and incubated overnight at either 37°C or 25°C as needed. The following day, the number of plaque-forming units (PFU) were counted to measure phage titer and assess infectivity through efficiency of plating (EOP). When individual plaques could not be discerned, a faint lysis zone across the drop area was considered to be 10 plaques.

### Liquid culture infection assays

Overnight cultures were diluted 1:100 in MMB with inducer and grown in tubes to OD_600_ 0.3 at 37°C. 180 μL of culture were transferred to Nunc™ MicroWell™ 96-Well, Nunclon Delta-Treated, Flat-Bottom Microplate (Thermo Fisher Scientific) and 20 μL of phages diluted in MMB were added to reach the appropriate MOI. The 96-well plate was then incubated at 37°C or 25°C with orbital shaking in an Infinite M200 plate reader (TECAN) for 6h. OD_600_ was measured every 10 min.

### Nuclease activity assays

Overnight cultures were diluted 1:100 in MMB with inducer and grown in tubes to OD_600_ 0.3 at 37°C. Cultures were split into two halves. In one half, phages were added to an MOI of 10. No phages were added to the other half as a control. At 0, 15 and 30 min post-infection at 25°C, samples were taken and plasmids and short DNA fragments were extracted using QIAprep Spin Miniprep Kit (Qiagen) according to manufacturer instructions. Extracted DNA was concentrated using a Concentrator plus (Eppendorf) with program V-AQ, 60°C, 15 min. The entire DNA extraction was then run on a 0.9% agarose gel containing ethidium bromide in Tris-acetate-EDTA buffer at 150V for 25 min and imaged with Gel Doc XR+ Imaging System (Bio-Rad) to observe DNA degradation.

### Phage burst size measurements

Overnight cultures were diluted 1:100 in MMB with inducer and grown in tubes to OD_600_ 0.3 at 37°C, shaking 200 rpm. SECphi18 phages were added to an MOI of 0.1 and infection proceeded at 25°C. As a control of initial phage titer, the same volume of phage was added to sterile MMB media and used as the titer of time 0 of infection. After 2h and 4h, corresponding to roughly one and two cycles of infection at 25°C for SECphi18, 1 mL of culture was collected, centrifuged for 3 min at 5000 *g* and filtered through a 0.2 μm filter. The phage titer was calculated as described above by doing a plaque assay infecting wild type *E. coli* MG1655.

### Growth curve experiments

Bacteria were streaked on LB 1.5% agar plates with the relevant antibiotics. One colony was resuspended in 50 μL MMB, then 5 μL of the resuspended culture were added to 195 μL MMB with antibiotic and appropriate inducer in Nunc™ MicroWell™ 96-Well, Nunclon Delta-Treated, Flat-Bottom Microplate (Thermo Fisher Scientific). Plates were then incubated at 37°C with orbital shaking in an Infinite M200 plate reader (TECAN) for 8 h and OD_600_ was measured every 10 min.

### Co-immunoprecipitation assays

Proteins were tagged with HA (YPYDVPDYA), 3xFLAG (DYKDHDGDYKDHDIDYKDDDDK) or TwinStrep (WSHPQFEKGGGSGGGSGGSAWSHPQFEK) tags as indicated in Supplementary Table S7, with a short linker between the tag and the protein of interest (SA for TwinStrep tag, SSG for HA and 3xFLAG). When co-expression of both proteins of interest was not possible due to toxicity, each protein was expressed in a separate strain and proteins were then mixed on beads, as described below.

Overnight cultures were diluted 1:100 in 50 mL MMB with antibiotics and grown for ∼2 h to OD_600_ ∼0.8 at 37°C, shaking 200 rpm. Then, the appropriate inducer was added and cells were incubated at 37°C for an additional hour. Cells were then pelleted and resuspended in 750 μL 1X Tween-Tris-buffered saline (TTBS, Biolab) with cOmplete™ ULTRA Tablets, Mini, EDTA-free, EASYpack Protease Inhibitor Cocktail (Roche) and kept on ice for the rest of the procedure. Cells were transferred to a Lysing Matrix B tube (MP biomedical) and broken down in a FastPrep-24™ Classic bead beating grinder (MP Biomedical) in two rounds of 40 s, 6 m/s shaking. Lysates were then centrifuged for 10 min at 12,000 *g* and 4°C and the supernatant was added to 25 μL Pierce™ Anti-HA Magnetic Beads (Thermo Fisher Scientific) washed in 1X TTBS buffer. Beads were incubated with the lysate for 1 h at 4°C with overhead shaking. Samples were placed on a magnet and the supernatant was removed. Beads were washed three times in 500 μL 1X cold TTBS buffer. When necessary, a second lysate containing the putative partner protein was added to the beads, incubated again for 1 h at 4°C and then washed again three times in 500 μL 1X cold TTBS. Proteins were then eluted by adding 25 μL 4X Bolt™ LDS Sample Buffer (Thermo Fisher Scientific) and heating for 5 min at 70°C to destroy the antibody. 10 μL of the supernatant were then used for western blotting.

### Western blotting

10 μL of eluted proteins were run on Bolt 4-12% Bis-Tris Plus Gels (Thermo Fisher Scientific) for 24 min at 200V in 1X Bolt™ MES SDS Running Buffer (Thermo Fisher Scientific). A PVDF membrane (Thermo Fisher Scientific) was activated in isopropanol for 90 s and then washed briefly in water. The proteins were transferred to the PVDF membrane in 1X Bolt™ transfer buffer (Thermo Fisher Scientific) for 1 h at 20 V in a Mini Blot module (Invitrogen). The membrane was then blocked for 30 min at room temperature or overnight at 4°C in 5% skim milk or 3% bovine serum albumin (specifically when detecting the TwinStrep tag) in 1X TTBS buffer. To detect the relevant proteins, the membrane was incubated with the appropriate primary antibody or Strep-Tactin^®^ HRP conjugate (IBA) diluted in 1X TTBS with 3% BSA for 1h at room temperature or overnight at 4°C. The membrane was then incubated with the secondary antibody prepared to appropriate dilution in 1X TTBS for 45 min at room temperature. No secondary antibody was used in the case of the HRP-coupled Strep-Tactin. Bands were stained using Luminata Forte Western HRP substrate ECL (Millipore) solution and imaged with ImageQuant™ LAS 4000 biomolecular imager (GE Healthcare). We used Spectra™ Multicolor Broad Range Protein Ladder (Thermo Fisher Scientific) as protein ladder. When necessary, the membranes were stripped using Restore PLUS Western Blot Stripping Buffer (Thermo Fisher Scientific) and probed with another set of primary/secondary antibodies, as detailed above. A list of all antibodies used in this study can be found in Supplementary Table S9.

### Isolation of phage escaper mutants

Phage mutants escaping the KpAvs2 defense system were isolated as described previously^24^. Briefly, 100 μL of bacterial cells expressing the KpAvs2 operon were grown in MMB supplemented with 0.2% arabinose to OD_600_ of 0.3 and then mixed with 100 μL SECphi18 phage lysate. After 10 minutes at room temperature, 5 mL pre-melted 0.5% MMB agar with 0.2% arabinose was added and the mixture was poured onto MMB 1.1% agar plates. The double layer plates were incubated overnight at room temperature and single plaques were picked into 90 μL phage buffer (50 mM Tris pH 7.4, 100 mM MgCl2, 10 mM NaCl). The collected phages were tested for their ability to infect both a control strain and an KpAvs2 operon-carrying strain and compared to parental SECphi18 phages by plaque assay, as described above. Phages which propagated better on the KpAvs2 operon-carrying strain than the parental phages were further amplified from a single plaque formed on the KpAvs2 operon-carrying strain as detailed below. Escaper 1372 was propagated in a liquid culture of bacteria expressing the KpAvs2 operon grown in 1 mL MMB with 0.2% arabinose to an OD_600_ of 0.3. The phages were incubated with the bacteria at 37°C 200 rpm shaking for 3 h, and then an additional 9 mL of bacterial culture grown to OD_600_ 0.3 in MMB with 0.2% arabinose was added, and incubated for another 3 hours (37°C 200 rpm). The lysate was then centrifuged at 3200 *g* for 10 min and the supernatant was filtered through a 0.2 μM filter to get rid of remaining bacteria. The other phages did not propagate as well in liquid culture, and thus the double layer plaque assay method was used for propagation. For this, the single plaque formed on the defense strain in the small drop plaque assay was picked into 100 μL phage buffer and mixed with 100 μL of cells grown in MMB with 0.2% arabinose to OD_600_ of 0.3. For escaper 1369, KpAvs2-operon expressing cells were used, and for escapers 1370, 1371, 1373, 1374, control cells expressing GFP were used. After incubation of the phages with the bacterial cells for 10 min at room temperature, 5 mL pre-melted 0.5% MMB agar with 0.2% arabinose was added and the mixture was poured onto MMB 1.1% agar plates. The double layer plates were incubated overnight at room temperature and 10^3^ -10^5^ PFUs were scraped into 5 mL of phage buffer. After 1 hour at room temperature, the phages were centrifuged at 3200 *g* for 10 min, and the supernatant was filtered through a 0.2 μM filter.

The obtained lysate was then used for DNA extraction and whole-genome sequencing. DNA was extracted from 500 μL of a high titer phage lysate (> 10^7^ PFU/mL). The phage lysate was treated with DNase-I (Merck cat #11284932001) added to a final concentration of 20 μg/mL and incubated at 37°C for 1 hour to remove bacterial DNA. DNA was then extracted using the DNeasy blood and tissue kit (Qiagen, cat #69504) starting from the Proteinase-K treatment step to lyse the phages. Libraries were prepared for Illumina sequencing using a modified Nextera protocol^35^. Reads were aligned to SECphi18 reference genome (NCBI accession NC_073071.1) and mutations compared to the reference genome were identified using breseq (version 0.29.0) with default parameters^36^. Only mutations that occurred in the isolated mutants, but not in the ancestor phage, were considered. A list of escaper phages obtained in this assay and the relevant mutations they carried can be found in Supplementary Table S3.

### Transformation efficiency assays

Plasmids carrying phage genes as indicated, or RFP as a control, were extracted from overnight cultures using QIAprep Spin Miniprep Kit (Qiagen) according to manufacturer instructions. DNA concentration was measured using Qubit® dsDNA HS Assay (Thermo Fisher Scientific).

Bacteria expressing KpAvs2 operon, either WT or lacking KpAvs2 C-terminal phage recognition domain, were grown to OD_600_ ∼0.3 in 4 mL MMB with ampicillin at 37°C, 200 rpm shaking. The samples were centrifuged 7 min at 2900 *g* and resuspended in 125 μL cold sterile Transformation and Storage Solution (TSS, LB broth with 10% PEG 3350 or 8000, 5% DMSO, and 40 mM MgCl2 at a final pH of 6.5) then split into 25 μL aliquots. 8 ng of each plasmid was added per tube and incubated 5 min on ice, then 5 min at room temperature and then again 5 min on ice^37^. Cells were allowed to recover in 1 mL MMB for 1 h at 37°C and 200 rpm shaking. Cells were then centrifuged for 4 min at 5000 *g* and serially diluted by 10-fold. 10 μL drops of each dilution were plated on LB+1.5% agar with ampicillin, kanamycin and glucose and incubated overnight at 37°C. The following day, colony-forming units (CFU) were counted as a proxy for transformation efficiency.

### Protein pulldown and LC-MS

150 mL of bacteria expressing either an HA-tagged, WT KpAvs2 protein, or a KpAvs2 protein lacking the C-terminal domain were grown to OD_600_ of 0.3 in the presence of 0.2 % arabinose at 37°C. Bacteria were then infected with SECphi18 at an MOI 5 at 25°C. 50 mL aliquots were collected before infection and 30 min post infection. The samples were immunoprecipitated using anti-HA antibodies as described above, except that the samples were eluted in 5% SDS in 100 mM Tris-HCl, pH 7.4 for 10 min at room temperature with occasional shaking rather than boiled in denaturing sample buffer. The eluted samples were then subjected to tryptic digestion using an S-Trap^38^. The resulting peptides were analyzed using nanoflow liquid chromatography (nanoAcquity) coupled to high resolution, high mass accuracy mass spectrometry (Q Exactive Plus). Each sample was analyzed on the instrument separately in a random order in discovery mode. Raw data was processed with MetaMorpheus v1.0.2. The data was searched against the *E. coli* K12 UniProt proteome database, the SECphi18 proteome, and additional proteins expressed on our plasmids and common lab protein contaminants, as well as default modifications. For KpAvs2, only the common sequence between the WT and deletion mutant was considered to avoid bias. Quantification was performed using the embedded FlasLFQ^39^ and protein inference^40^ algorithms. The LFQ (Label-Free Quantification) intensities were calculated and used for further calculations using Perseus v1.6.2.3. Decoy hits and contaminants were filtered out. The LFQ intensities were log transformed and only proteins that had at least two valid values in at least one experimental group were kept. The remaining missing values were imputed by a random low range normal distribution. A student’s t-test was performed to identify differentially represented proteins. The mass spectrometry proteomics data was deposited to the ProteomeXchange Consortium^41^ via the PRIDE^42^ partner repository with the dataset identifier PXD048766.

### Structural analyses and remote homology detection

Structural predictions of proteins and complexes were performed using AlphaFold2^31^ and AlphaFold-Multimer^21^, respectively, version 2.3.1 with default parameters. Five predictions for complexes or one prediction for single chain proteins were computed for five models for each structure, and the best scoring model was considered further. Structures were visualized using the PyMOL Molecular Graphics System, Version 2.5.2 (Schrödinger, LLC). When applicable, proteins were structurally aligned using the cealign function of PyMOL. To predict the list of residues interacting between two protein chains, the Residue Interaction Network Generator (https://ring.biocomputingup.it/submit)^29^ was used.

HHpred^19,20^ was used to annotate genes for which automatic sequence-based annotation did not retrieve annotations (toolkit.tuebingen.mpg.de/tools/hhpred).

### Identification of KpAvs2 and Ksap1 homologs

Homologs of KpAvs2 were identified from both isolates and metagenomes samples from the IMG^22^ database in April 2020 as described previously^43^. The genomic neighborhood of each member of the KpAvs2 cluster was examined, and only homologs that were found in a conserved three-gene operon were retained.

Homologs of Ksap1 were identified by blast search on the National Center for Biotechnology Information (NCBI, https://blast.ncbi.nlm.nih.gov/Blast.cgi)^44,45^ website. Both blastP and tblastN were used with default parameters. Ksap1 homologs that were not annotated in their genome of origin were identified manually by examining the intergenic region in the syntenic loci in all genomes of *Dhillonviruses* present in the NCBI Taxonomy browser that failed to show a hit in either search for Ksap1, or showed a hit only to a part of Ksap1. The genome of SECphi6 (NCBI accession CADCZA000000000.2) was similarly searched. A list of all identified homologs is found in Supplementary Table S6. All identified homologs were manually curated, aligned with Clustal Omega^46^ and a web logo was created using the online web logo tool, version 2.8.2 (weblogo.berkeley.edu/logo.cgi)^47,48^. An AlphaFold2-predicted protein structure of Ksap1 was used to search for structural homologs using the Foldseek search^30^ website (search.foldseek.com/search), using default databases and 3Di/AA mode.

## Acknowledgements

We thank members of the Sorek laboratory for comments on earlier versions of this manuscript and in particular Jens Hör for helpful discussions on pulldown experiments design. We also thank Alon Savidor and Meital Kupervaser at the De Botton Protein Profiling Institute at the Weizmann Institute for help with mass spectrometry data generation and analysis, and the members of the Crown Genomics institute of the Nancy and Stephen Grand Israel National Center for Personalized Medicine for help with Sanger DNA sequencing. R.S. was supported, in part, by the European Research Council (grant no. ERC-AdG GA 101018520), Israel Science Foundation (MAPATS Grant 2720/22), the Deutsche Forschungsgemeinschaft (SPP 2330, Grant 464312965), the Ernest and Bonnie Beutler Research Program of Excellence in Genomic Medicine, Dr. Barry Sherman Institute for Medicinal Chemistry, Miel de Botton, the Andre Deloro Prize, and the Knell Family Center for Microbiology. N. B. was supported by a postdoctoral grant from the Azrieli foundation and a fellowship from the Deans of the Faculty of Weizmann Institute of Science.

## Supplementary Materials

### Supplementary Figures

**Supplementary Figure S1.**
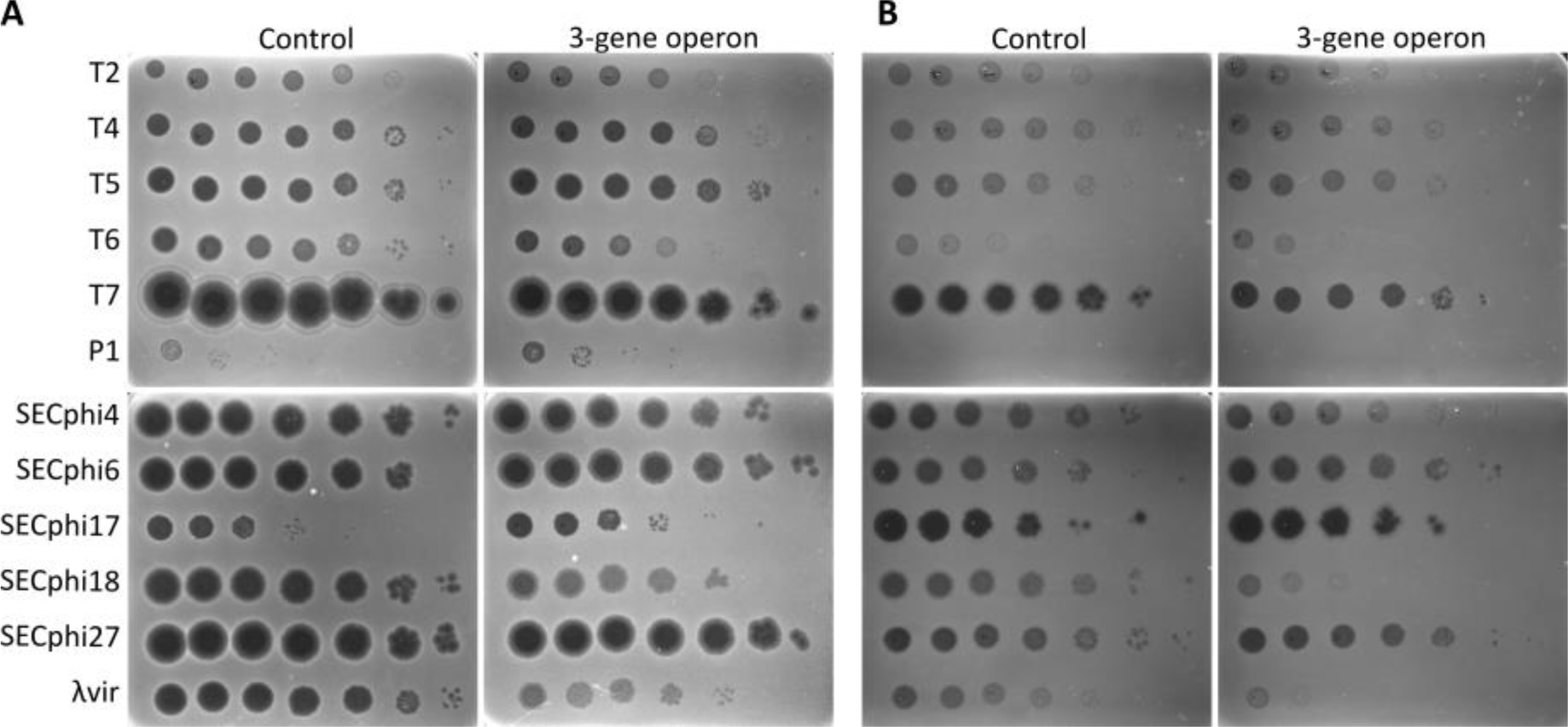
KpAvs2 operon-mediated defense against *E. coli* phages. A representative plaque assay showing efficiency of plating of different phages on a lawn of cells expressing the KpAvs2 operon, or control cells expressing GFP instead. Infection was performed at **A.** 37°C or **B.** 25°C. Quantification for phages for which defense was observed is presented in Figure 1C.

**Supplementary Figure S2.**
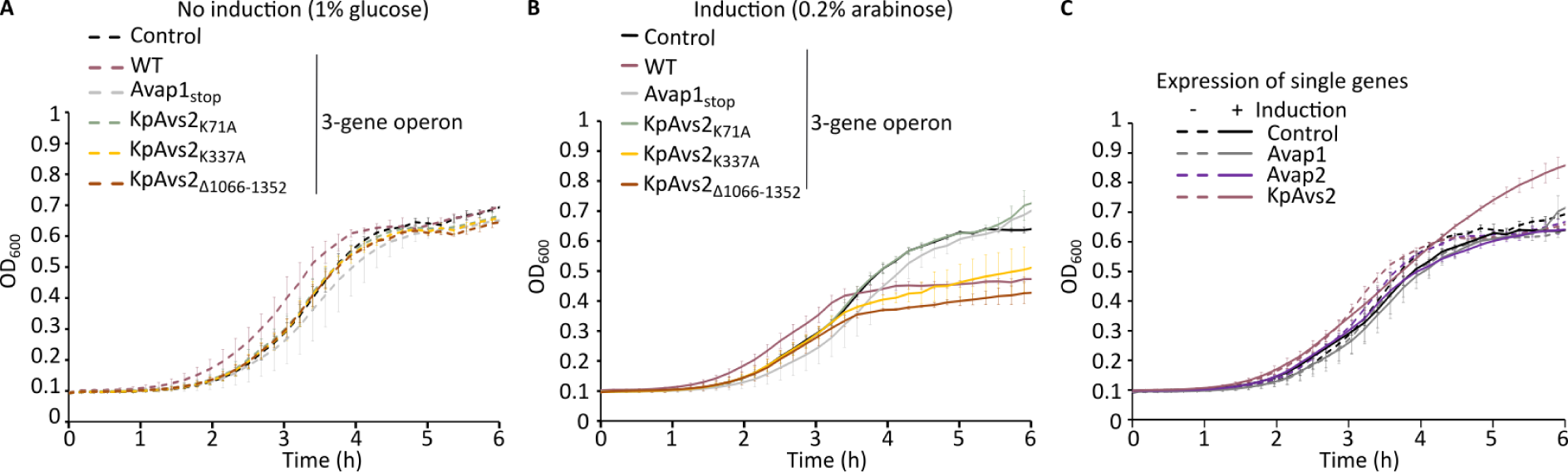
Growth of KpAvs2 operon mutants in liquid culture. Growth curves in liquid culture of *E. coli* cells expressing the KpAvs2 operon with the indicated mutation, shown here as no-infection controls for the experiments presented in Figure 1D. *E. coli* cells with a plasmid carrying GFP were used as control. Bacteria were grown at 37°C. Data represent the average of three replicates, error bars represent standard deviation. **A.** Growth in absence of induction. 1% glucose was used to repress expression. **B.** Growth when expression is induced using 0.2% arabinose. **C.** Growth in the presence of 1% glucose (-) or 0.2% arabinose (+) of strains expressing either GFP (control) or single proteins of the KpAvs2 operon.

**Supplementary Figure S3.**
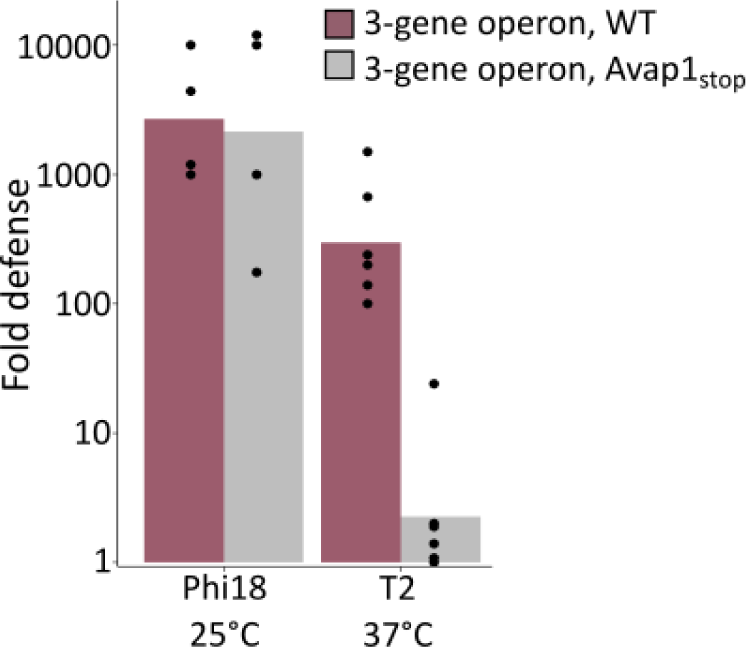
Avap1 is necessary for defense against T2. *E. coli* cells expressing the WT KpAvs2 operon, or an operon with a premature stop codon in Avap1, were infected by SECphi18 and T2. Fold defense was calculated as the ratio between the efficiency of plating of phages on control cells expressing GFP divided by the efficiency of plating observed for the indicated strain. Bar graph represents the average of 4-6 independent replicates, with individual data points overlaid.

**Supplementary Figure S4.**
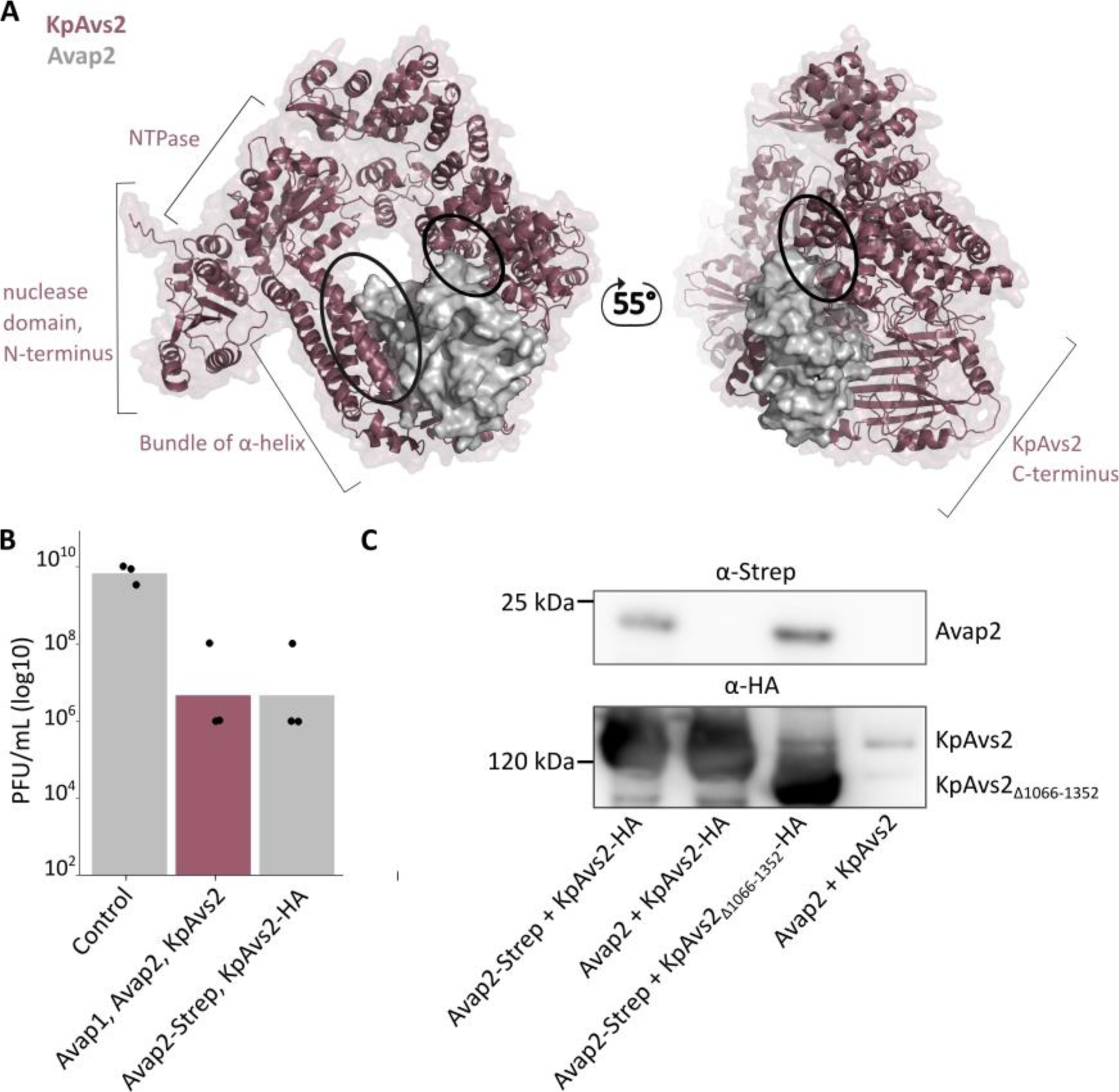
Avap2 binds KpAvs2. **A.** Interactions between Avap2 (in grey) and KpAvs2 (in purple) predicted by AlphaFold-Multimer^21^. Model confidence score: 0.809. Dark circles highlight the two predicted zones of physical binding between the two proteins. **B.** Tagging of KpAvs2 and Avap2 does not interfere with the anti-phage properties of the system. Efficiency of plating of SECphi18 phage on *E. coli* cells expressing GFP (control), the KpAvs2 operon (Avap1 Avap2 KpAvs2) or a TwinStrep-tagged Avap2 and an HA-tagged KpAvs2 (Avap2-Strep, KpAvs2-HA). Infection proceeded at 25°C. Data represent plaque-forming units (PFU) per mL. Bar graph represents the average of 3 independent replicates, with individual data points overlaid. **C.** Co-immunoprecipitation of HA-tagged KpAvs2 and TwinStrep-tagged Avap2. α-HA beads were used to co-immunoprecipitate HA-tagged KpAvs2, and the interacting Avap2 was detected by western blot against TwinStrep tag. A western against HA was performed on the same blot to control for efficiency of pulldown. KpAvs2 lacking the C-terminus was used as a control (KpAvs2_Δ1066-1352_). Controls expressing the untagged proteins were also used, as indicated.

**Supplementary Figure S5.**
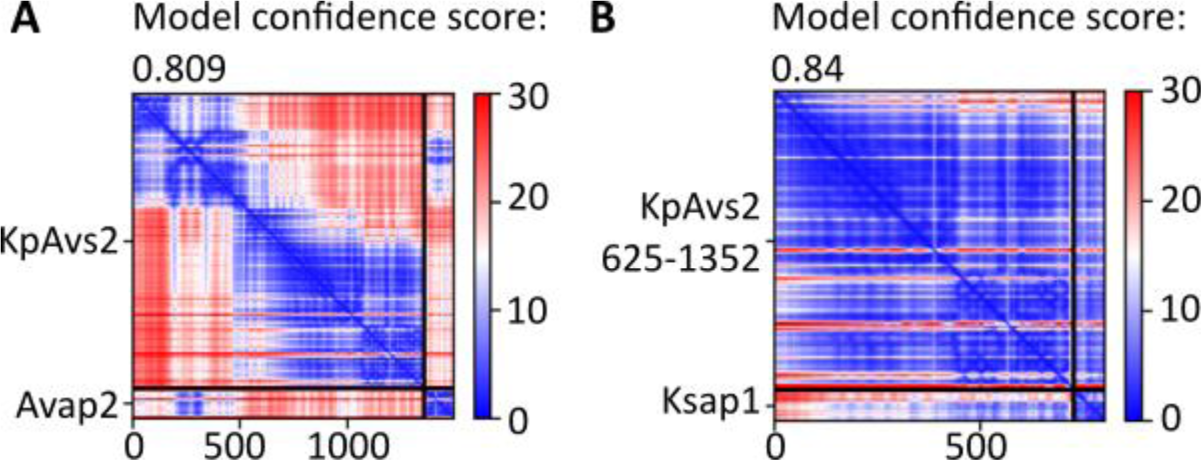
AlphaFold-Multimer predictions. **A.** Predicted aligned error (PAE)^31^ plot of the AlphaFold-Multimer^21^ prediction shown in Supplementary figure S4A. **B.** PAE plot of the structure shown in Figure 3C.

**Supplementary Figure S6.**
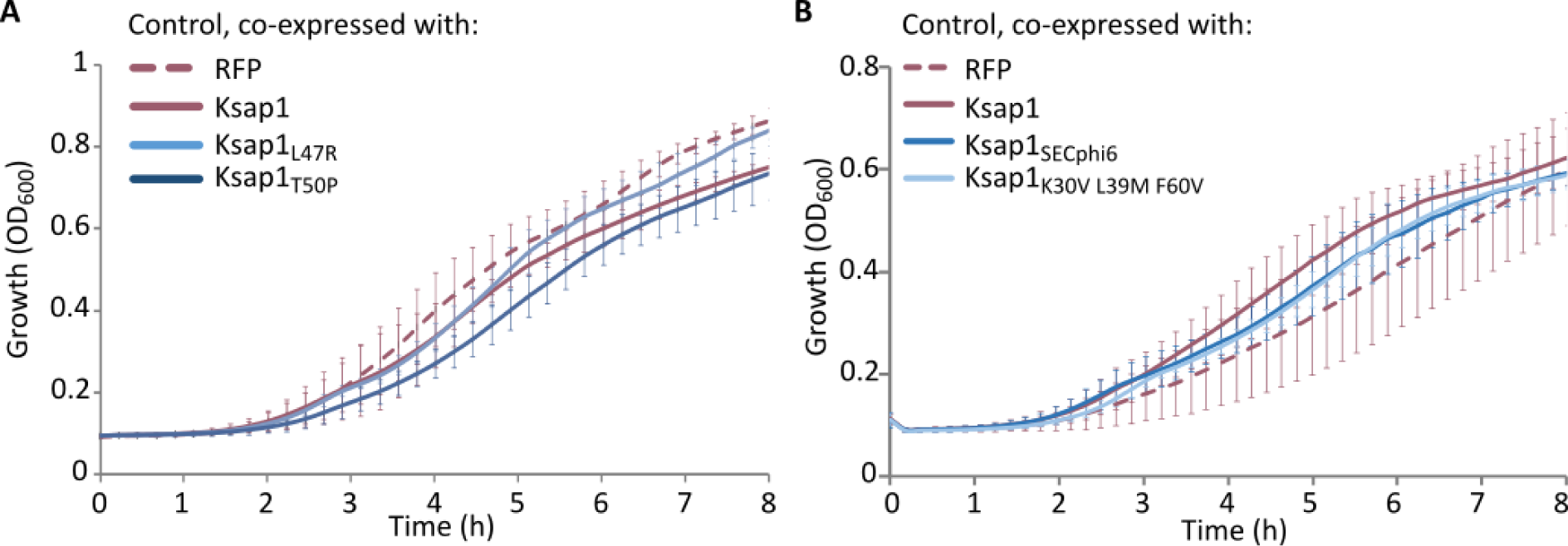
Control for of Ksap1 and KpAvs2 co-expression assays. Growth curves in liquid media of *E. coli* cells co-expressing GFP and Ksap1 variants as indicated. A plasmid carrying RFP instead of Ksap1 was used as control. Data represent the average of six replicates, error bars represent standard deviation. **A.** Related to Figure 3G. **B.** Related to Figure 4B.

**Supplementary Figure S7.**
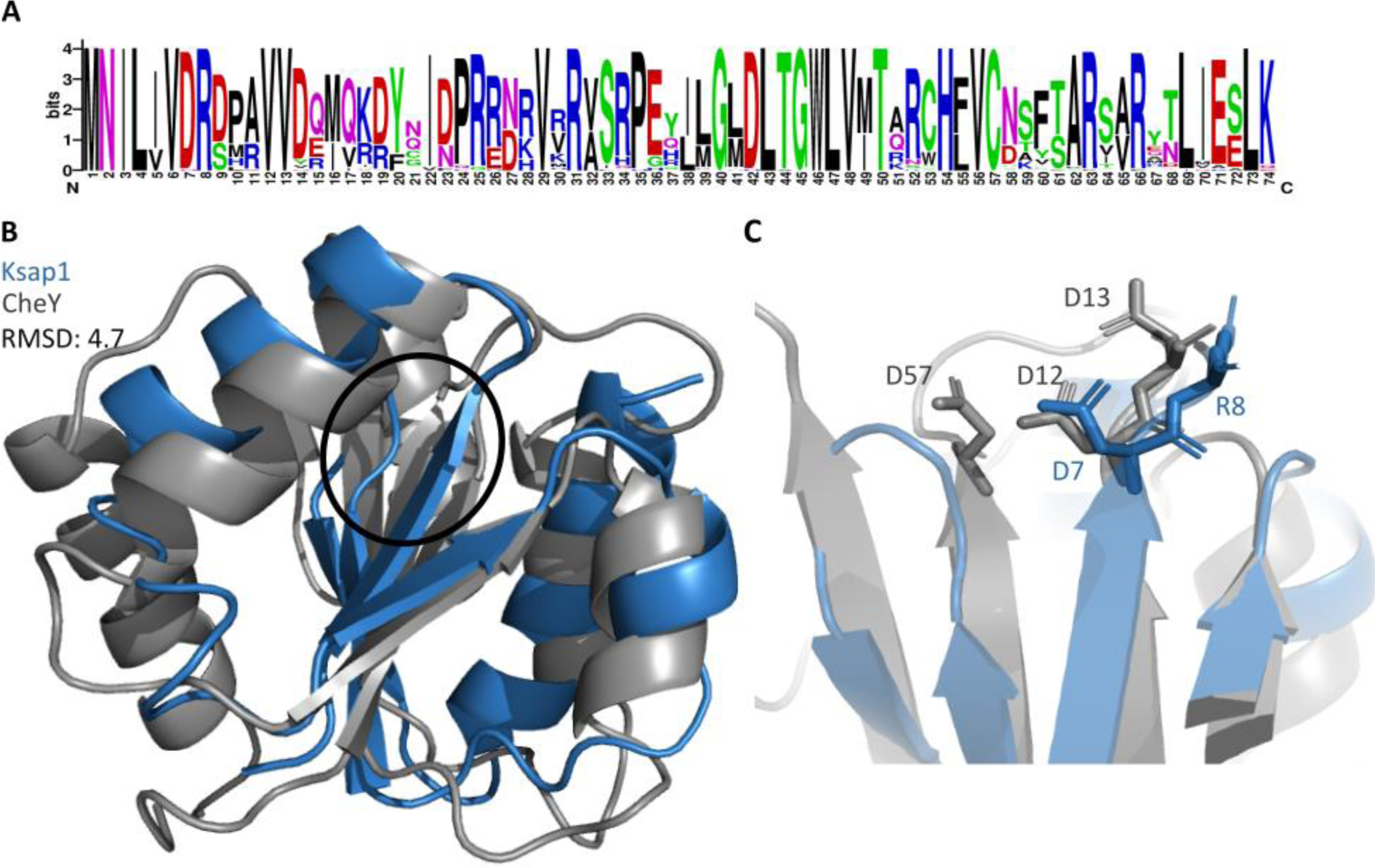
Ksap1 homologs. **A.** Multiple sequence alignment of Ksap1 homologs identified in NCBI. Alignment was performed with Clustal Omega^46^ and represented as a Web Logo^47^. **B.** Structural alignment between the crystal structure of a CheY response regulator domain (PDB 2CHF, grey)^49^ and the AlphaFold2-predicted structure of Ksap1 (blue). A black circle represents the area that is shown in more detail in panel C. **C.** A close-up on three aspartic acid residues in CheY that are essential for phosphorylation^33^ (grey), and the corresponding residues in Ksap1 (blue).

### Supplementary Tables

Supplementary Table S1. KpAvs2 homologs identified as part of a conserved three-gene locus.

Supplementary Table S2. Proteins identified by mass spectrometry after KpAvs2 pulldown.

Supplementary Table S3. Mutations identified in escaper phages genomes.

Supplementary Table S4. Predicted interactions between KpAvs2 and Ksap1.

Supplementary Table S5. Predicted interactions between KpAvs2 and Ksap1_SECphi6_.

Supplementary Table S6. Ksap1 homologs.

Supplementary Table S7. Strains, plasmids and phages used in this study.

Supplementary Table S8. Primers used in this study.

Supplementary Table S9. Antibodies used in this study.

